# Menin maintains enhancer-promoter interactions in a leukemia-specific manner

**DOI:** 10.64898/2026.01.16.698179

**Authors:** Vassilena Sharlandjieva, Catherine Chahrour, Frederik H. Lassen, Joseph C. Hamley, Andreas Damianou, Nicholas Denny, Alastair L. Smith, Svenja S. Hester, Iolanda Vendrell, Ronald W. Stam, Marina Konopleva, Anindita Roy, James O. J. Davies, Nicholas T. Crump, Benedikt M. Kessler, Thomas A. Milne

## Abstract

Inhibition of the protein-protein interaction between Mixed Lineage Leukemia (MLL) and Menin is a promising therapy for both high-risk *MLL*-rearranged and *NPM1*-mutant (NPM1c) acute leukemias, yet the mechanistic basis of this dependency in distinct contexts remains unclear. By comparing the transcriptional responses of MLL::AF4 and NPM1c leukemia models to Menin inhibition, we find broad, acute transcriptional dysregulation in MLL::AF4 cells, but minor transcriptional consequences in NPM1c cells, despite similarities in Menin promoter occupancy. Using high-resolution Micro Capture-C, we discover that Menin drives enhancer activity and maintains enhancer-promoter contacts in MLL::AF4 cells but not in NPM1c cells. Crucially, Menin is also essential for patient-specific enhancer function in primary *MLL*-rearranged leukemia samples. Proteomic analysis further demonstrates that Menin associates with distinct transcriptional and elongation complexes in MLL::AF4 compared to NPM1c cells, supporting a context-dependent mechanism of action. Together, these findings establish that Menin is not a uniform transcriptional cofactor, but a context-dependent regulator of enhancer connectivity, and identifies enhancer-promoter architecture as a selective vulnerability in MLL-rearranged leukemia.

## Introduction

Chromatin is the DNA/protein complex in the cell that influences both transcriptional activity and cell identity.^1^ In leukemia, frequent mutations in chromatin-modifying proteins and transcription factors trigger oncogenic transcription, often hijacking endogenous machinery, which leads to the expansion of immature progenitor cells.^2,3^ As a result, epigenetic enzymes, including transcriptional repressors (EZH2, HDAC, LSD1) and activators (P300, bromodomain proteins, DOT1L),^4–6^ have become promising targets in hematological malignancies.

Menin is a chromatin protein that has emerged as a key vulnerability in high-risk leukemia. Menin is a component of multiple complexes,^7–10^ but its best studied role is as a member of the Mixed Lineage Leukemia complex (MLL1 or MLL, also known as KMT2A), where it interacts directly with the N-terminus of MLL. Rearrangements of the *MLL* gene (*MLL*r) give rise to oncogenic fusion proteins sufficient to drive leukemia, and account for approximately 10% of acute leukemias overall but up to 90% of infant B acute lymphoblastic leukemia (ALL).^11–13^ Patients with *MLL* rearrangements have a worse prognosis and a poor response to conventional chemotherapy,^14^ highlighting an unmet clinical need. MLL fusion proteins (MLL-FPs), most commonly MLL::AF4 in infants and MLL-AF9 in adults,^15^ assemble a large protein complex, including Menin, which activates gene expression through both epigenetic and transcription elongation mechanisms.^16–18^ Menin is essential for leukemia initiation in the presence of MLL-FPs, facilitating oncogenic transcription by stabilizing MLL-FPs at hematopoietic genes, such as *MEIS1.*^19^

Menin inhibitors disrupt the Menin-MLL/MLL-FP interaction and initially aimed to target *MLL*r leukemia.^20^ However, they have also been found to be effective in acute myeloid leukemia (AML) with *NPM1* mutations.^21–25^ Affecting 30% of patients, *NPM1* mutations are the most common genetic lesion in AML,^26^ meaning that Menin inhibition (MENi) has the potential to benefit a large group of patients. *NPM1* mutations are commonly frameshifts in the C-terminus that disrupt a nucleolar localisation signal.^27–30^ They have a dual effect on the typically nucleolar nucleophosmin protein: a portion of the mutant protein (“NPM1c”) aberrantly localizes to the cytoplasm, while a small proportion in the nucleus can function as a transcription activator by directly binding at a subset of Menin-MLL target genes.^21,24^ NPM1c is associated with expression of the *HOXA/B* cluster and their co-factor *MEIS1,* similar to MLLr leukemias.^31,32^

Menin inhibitors cause the loss of Menin binding at chromatin and downregulation of key MLL-FP and NPM1c targets.^22,33^ However, as Menin is part of distinct complexes in *MLLr* and NPM1c leukemias, it is unclear if Menin has direct role in driving transcription in these distinct leukemia subtypes. Both MLL-FPs and NPM1c are transcriptional activators, binding at active promoters and spreading into gene bodies.^24,34^ In addition to promoters, MLL-FPs have also been shown to bind enhancers,^35–37^ non-coding regulatory elements that modulate gene transcription from a distance.^38^ The resulting aberrant enhancer activation is a key mechanism driving oncogenic gene expression programmes in *MLL*r leukemia.^36,39,40^ While this implicates Menin in the maintenance of enhancer activity in the presence of MLL-FPs, Menin also binds a subset of enhancers in ER+ breast cancer cells with wildtype *MLL*, suggesting a possible role for Menin in driving enhancer function more generally.^41^ NPM1c binding outside promoters has also been reported,^24^ but whether it is functionally maintaining enhancer activity, and whether it requires Menin for this, remains unknown. Better understanding the role of Menin in transcription in different contexts could inform our interpretation of patient response, relapse and resistance in ongoing clinical trials (summarized in^42^ and^43^).

In this paper, we show that MENi triggers large scale, acute transcriptional dysregulation in MLL::AF4 ALL cells but not in NPM1c AML cells. This difference is associated with Menin binding at intragenic enhancers in the presence of MLL::AF4, where it facilitates enhancer-promoter contacts at key oncogenes, an activity absent in NPM1c cells. We also show a greater enrichment of transcription regulators and elongation factors in the Menin interactome in MLL::AF4 cells compared to NPM1c cells, suggesting a more central role for Menin in gene activation. Together, our characterization of Menin activity in distinct leukemia subtypes demonstrates that the mechanism by which MENi disrupts oncogenic transcription is unique to context-specific Menin function.

## Results

### Menin inhibition has distinct transcriptional effects in MLL::AF4 and NPM1c cells

While current models for the therapeutic function of MENi state that destabilized MLL-FP or NPM1c binding trigger decreased gene expression,^44^ most studies of the effects of MENi report transcriptional changes after multi-day treatments and so early, direct effects on Menin-bound genes in different contexts are unclear.^25,33^ We sought to compare the effects of MENi on nascent transcription at an early timepoint, using SEM and OCI-AML3 cells as models of MLL::AF4 ALL and NPM1c AML respectively. As a reference to a model of insensitivity to MENi, we used RCH-ACV cells, an E2A-PBX1-driven ALL cell line. Following treatment with the Menin inhibitor VTP50469 (250 nM) for 24 hours, Menin binding was depleted from transcription start sites (TSS) genome-wide in each cell line (Figure S1A), regardless of inhibitor sensitivity status. Transient Transcriptome RNA-seq (TT-seq) at this timepoint revealed extensive transcriptional changes in SEM (MLL::AF4) cells, and surprisingly few changes in OCI-AML3 (NPM1c) cells (Figure 1A). MENi resulted in 1874 differentially expressed genes in SEM cells, the majority (1058) of which were downregulated. They included MLL::AF4 targets such as *FLT3, MEF2C* and *JMJD1C,^33^* but not *HOXA9* or *MEIS1*. In contrast, OCI-AML3 (NPM1c) cells had 103 differentially expressed genes of which 72 were downregulated, including *MEIS1*, which is considered the critical Menin-dependent vulnerability in these cells.^21,23,25^

**Figure 1.:**
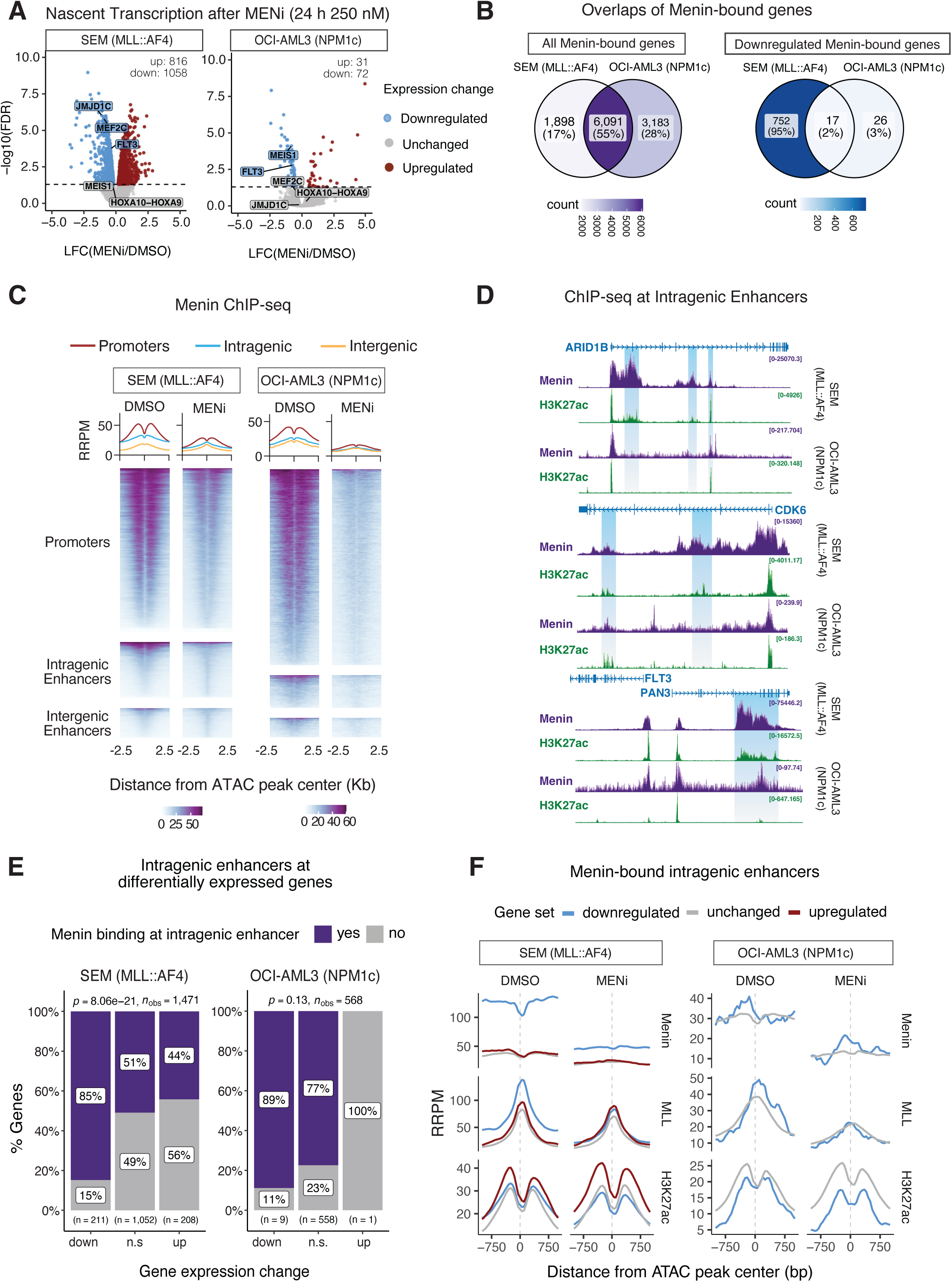
Differential effects of Menin inhibition on transcription in MLL::AF4 ALL and NPM1-mutant AML. (A) Volcano plots of transient transcriptome RNA-seq after 24 h of treatment with DMSO or 250 nM Menin inhibitor VTP50469 (MENi). (B) Overlap of all Menin-bound genes (left) and downregulated Menin-bound genes only (right) in SEM and OCI-AML3 cells. (C) Heatmaps of Menin ChIP-seq at promoters and enhancers in SEM and OCI-AML3 cells. Regions are sorted by H3K27ac signal and centered on ATAC peaks. Metaplots display the mean. (B) Examples of intragenic Menin-bound enhancers in the gene bodies of *ARID1B*, *CDK6* and *PAN3*. (D) Association between Menin binding at intragenic enhancers and altered gene expression from the nearest TSS following 24 h of MENi. *p*-values are reported from a Pearson chi-square test. (E) Mean Menin, MLL/MLL::AF4, and H3K27ac at intragenic enhancers annotated to differentially expressed genes by nearest TSS after 24 h of MENi in SEM and OCI-AML3 cells.

This contrast in transcriptional consequences following MENi was correlated with effects on cell fitness at 48 h MENi. Using CellTiterGlo assays, we observed that SEM cells were highly sensitive, with an IC50 of 45.5 nM, whereas OCI-AML3 cells did not reach IC50 even at the highest concentration of inhibitor (62.5 µM) tested, similar to the insensitive RCH-ACV cells. (Figure S1B). In contrast, colony formation assays over a two-week period showed that both SEM and OCI-AML3 colony formation was inhibited completely with MENi (Figure S1C).

To understand why Menin loss has broad and immediate transcriptional consequences in SEM (MLL::AF4) cells yet highly restricted effects in OCI-AML3 (NPM1c) cells, we compared Menin occupancy at the promoters of differentially expressed genes in each cell line. Curiously, we found a significant association between Menin promoter binding and gene downregulation in SEM (*p* = 1.52e-45, chi-square test) but not in OCI-AML3 (*p* = 0.13) (Figure S1D). Downregulated genes in SEM cells also had higher average Menin enrichment compared to upregulated or unchanged genes (Figure S1E). Yet, when we intersected Menin-bound promoters in the two cell lines, we found that the majority (6,091) were shared between cell lines (Figure 1B, left), while down-regulated Menin targets were predominantly unique to SEM cells (Figure 1C, right). This indicated that common Menin occupancy at promoters was not predictive of transcriptional sensitivity to MENi in SEM and OCI-AML3 cells.

Menin is thought to stabilize MLL::AF4 and NPM1c chromatin binding and both these oncoproteins have been shown to activate genes by recruiting RNA polymerase II and transcription elongation factors.^24,45,46^ Previous work has identified 2,597 unique genes bound by MLL::AF4 in SEM cells,^34^ whereas only 44 genes are NPM1c-bound in OCI-AML3 cells.^21^ It is therefore possible that the context-dependent regulation of genes by these oncoproteins determines the scale of transcription dysregulation in response to MENi. However, MENi disrupts MLL::AF4 binding at only a subset of its target promoters,^33^ and indeed we also saw that after 24 h MENi, only 16.1% (176/1058) of downregulated genes in SEM cells had a concordant decrease in MLL::AF4 promoter occupancy (Figure S1F, G). Similarly, 8/72 down-regulated genes in OCI-AML3 cells had decreased NPM1c binding. Taken together, our data indicated that neither high Menin occupancy at promoters nor loss of MLL::AF4 binding can fully explain the large number of dysregulated genes in SEM (MLL::AF4) cells following MENi.

### Menin binds enhancer elements

To further investigate why Menin has a more dramatic and immediate effect on transcription in SEM cells and not OCI-AML3 cells, we probed for the presence of Menin binding outside of gene promoters. We annotated Menin ChIP-seq peaks to promoters, intragenic enhancers and intergenic enhancers, and linked them to genes using the nearest TSS. This confirmed Menin binding at a subset of enhancers in both SEM and OCI-AML3 cells, although less frequently than its known enrichment at promoters (Figure 1C). For example, we saw Menin-bound domains overlapping with H3K27ac in the gene bodies of *ARID1B*, *CDK6*, and *PAN3,* all of which were larger in SEM (MLL::AF4) cells relative to OCI-AML3 cells (NPM1c) (Figure 1D). Using publicly available AutoCUT&Tag^47^ and ChIP-seq data, we also confirmed Menin enrichment outside promoters in additional *MLL*r cell lines (Figure S2A, B), as well as Menin binding at intragenic enhancers in primary patient cells which we profiled using small cell number ChIPmentation^48^ (Figure S2C), indicating that this phenomenon extends beyond the SEM cell model.

We found that Menin-bound intragenic enhancers were specifically enriched in the proximity of downregulated genes in SEM cells but not OCI-AML3 cells (Figure 1E). This link was stronger for intragenic enhancers compared to intergenic enhancers (Figure S2D); we also identified more total genes linked to Menin-bound intragenic enhancers (807, 55% of genes with an intragenic enhancer) versus intergenic enhancers (220, 35% of genes with an intergenic enhancer) in SEM cells. This observation was in line with previous work showing that Menin can spread into gene bodies in the presence of MLL::AF4.^34^ Intragenic Menin-bound enhancers linked to downregulated genes had higher levels of Menin and MLL/MLL::AF4 in SEM cells, and to a lesser extent in OCI-AML3 cells (Figure 1F). Upon MENi, intragenic enhancers had reduced MLL::AF4 / MLL binding and, more subtly, H3K27ac. A similar effect was observed at Menin-bound intergenic enhancers (Figure S2E). Given the unique enrichment of Menin-bound putative intragenic enhancers in SEM cells, we sought to further characterise the effects of Menin inhibition on enhancer activity in our cell line models.

### Menin regulates enhancer activity in MLL::AF4 cells

A key attribute of enhancers is physical proximity with their target promoters in the 3D organisation of the genome.^49–51^ As a result, annotating enhancers to the nearest TSS may not reveal the true target promoter; instead, high-resolution Chromatin Confirmation Capture techniques are needed to link enhancers and promoters. We have previously shown using Capture-C that enhancers occupied by the MLL::AF4 complex lose promoter contact upon knock-down or enzymatic inhibition of complex components.^36,37^ To test whether Menin binding also maintains enhancer-promoter contacts, we used the higher-resolution Micro Capture-C (MCC)^52^ to identify enhancer contacts for key gene promoters in SEM and OCI-AML3 cells (Figure 2A). Our panel of promoter viewpoints included key MLL::AF4 and NPM1c targets, genes sensitive to MENi, and genes linked to Menin-bound enhancers using nearest TSS annotation in both cell lines (Supplementary File 1, Figure S3A).

**Figure 2.:**
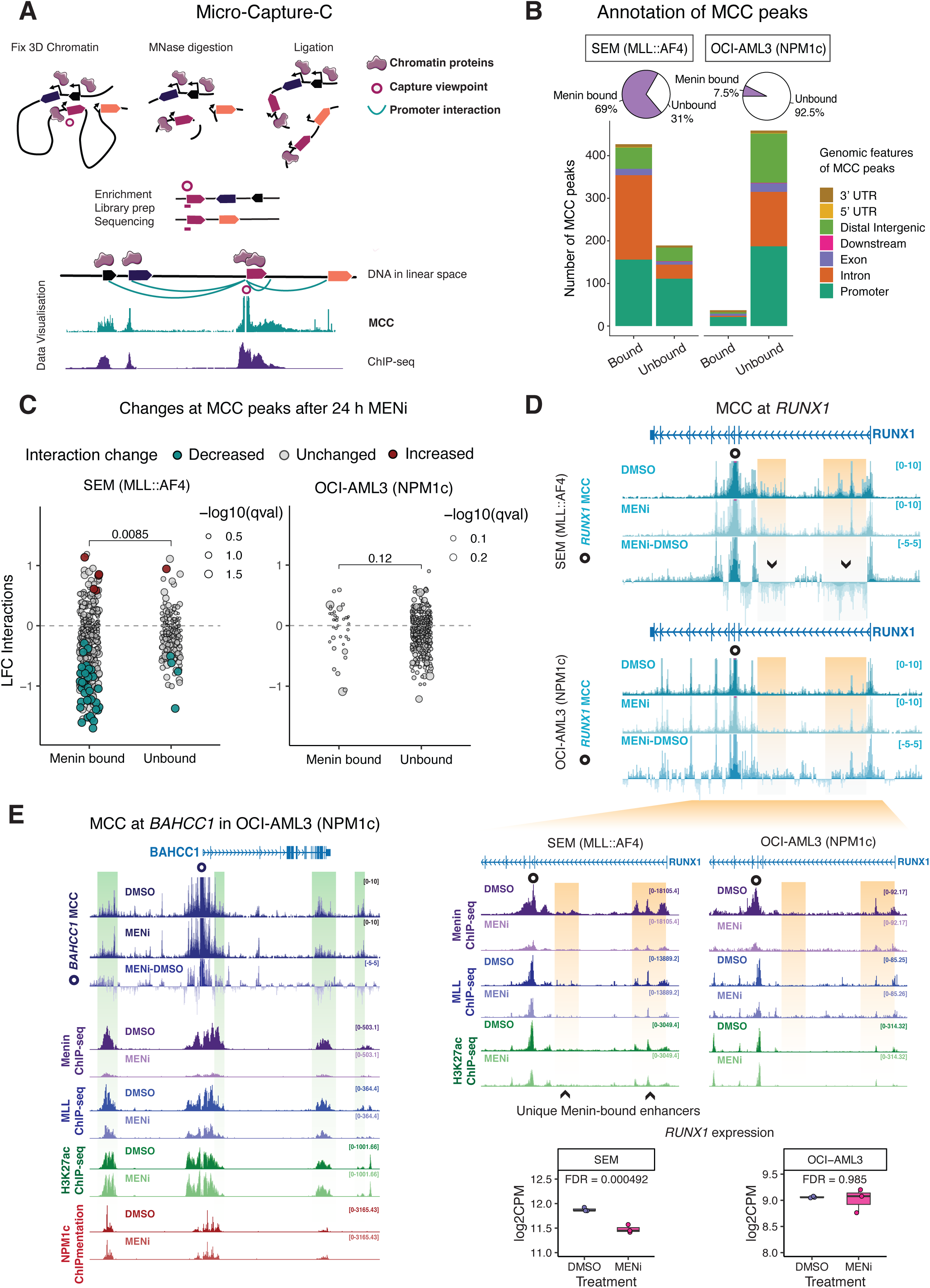
Menin inhibition disrupts enhancer-promoter contacts in MLL::AF4 cells but not NPM1c cells. (A) Schematic representation of Micro Capture-C data. (B) Annotation of MCC peaks (enhancer-promoter contacts), grouped by the presence of Menin binding, in SEM and OCI-AML3 cells. (C) Changes in mean unique ligation junction counts at MCC peaks in SEM (left) and OCI-AML3 (right) cells treated with MENi (250 nM, 24 h), categorised by Menin binding. Colored data points represent significantly altered contacts (Mann-Whitney U, FDR-adjusted q-value < 0.05). *p*-values comparing bound and unbound sites are reported from a Wilcoxon test. (D) Visualization of MCC (top, mean+whiskers signal for DMSO and MENi), ChIP-seq (middle, mean signal), and gene expression (bottom, TT-seq data) at the *RUNX1* locus in SEM and OCI-AML3 cells. The MENi-DMSO subtraction track shows the difference in MCC signal (mean with whiskers) between conditions. Menin-bound enhancers in SEM cells are highlighted in yellow. (E) Visualization of MCC (top, mean+whiskers) and ChIP-seq (bottom, mean signal) at the *BAHCC1* locus in OCI-AML3 cells. Menin-bound enhancers are highlighted in green.

We filtered MCC peaks (representing regions in contact with the sampled gene promoters) to those overlapping with H3K27ac to define putative functional promoter interactions. Menin was enriched at 69% of promoter interactions for 43 genes analysed in SEM (MLL::AF4) cells, but only 7.5% of interaction sites for 42 genes analysed in OCI-AML3 (NPM1c) cells (Figure 2B). The majority (46%) of Menin-bound enhancers in SEM cells were within introns, followed by MCC peaks (contacts) in other promoters (37%). In contrast, Menin was primarily enriched in other promoter regions (57%) in contact with the OCI-AML3 promoter viewpoints. This contrast emphasized that Menin function is primarily restricted to key promoters in OCI-AML3 cells, whereas intragenic spreading^34^ in SEM cells can be linked to enhancer activity.

Clustering promoter contact sites (MCC peaks) based on chromatin features revealed that Menin-bound enhancer-promoter contacts also had higher MLL/MLL::AF4 and H3K27ac in both cell lines, as well as NPM1c enrichment OCI-AML3 (Figure S3B). Upon MENi for 24 hours, enhancers with the highest levels of Menin (cluster 1, Figure S3B) also lost MLL::AF4 binding and H3K27ac in SEM cells. In contrast, in OCI-AML3 cells, H3K27ac and NPM1c remained unaffected at the Menin-enriched contacts, suggesting a unique sensitivity to MENi in SEM cells.

We next asked whether MENi for 24 hours affected the ability of enhancer loci to interact with target promoters. To quantify changes in enhancer-promoter interactions upon MENi across all the sampled viewpoints, we calculated the change in the mean count of unique ligation junctions at each interaction site. In SEM cells, 49 interaction sites had a significant change in interaction frequency (Mann Whitney U test, FDR q-value < 0.05); of the decreased interactions, 90% (39/43) overlapped with Menin binding (Figure 2C). The decrease in promoter interactions with Menin-bound enhancers was significantly lower than sites lacking Menin binding (Wilcoxon test *p*-value < 0.01), pointing to a specific Menin inhibitor-driven effect. Conversely, there were no significant changes in interactions in OCI-AML3 cells, including at Menin-bound sites, suggesting that Menin is dispensable for enhancer-promoter contacts in these cells.

The *RUNX1* locus illustrated the differential effects of MENi on enhancer activity (Figure 2D). In both SEM and OCI-AML3 cells, the major transcript isoform of *RUNX1* initiates at intron 3 (Figure S3C), so we used this site for the MCC capture enrichment. This region contacts the *RUNX1* promoter and several regions in intron 2 (Figure 2D, top, yellow highlights). In SEM cells, Menin inhibition leads to a decrease in interaction frequency with enhancers in intron 2, which we did not observe in OCI-AML3 cells. Sites of decreased interaction had higher Menin enrichment in SEM cells, indicating an SEM-specific Menin-bound enhancer cluster (Figure 2D, bottom, yellow highlights). The collapse of the intragenic Menin-bound enhancers in SEM cells correlates with decreased *RUNX1* expression following MENi (Figure 2D, bottom), whereas expression was unaffected in OCI-AML3 cells.

In OCI-AML3 cells, the NPM1c target gene *BAHCC1* has three Menin-bound enhancer regions (Figure 2E). At two of them, we observed a decreased interaction frequency on average (log2 fold change < 0), but this was not statistically significant (FDR > 0.05). H3K27ac, NPM1c binding, and gene expression were also unaffected despite Menin loss (Figure 2E, Figure S3D). This suggests that while Menin may support enhancer-promoter contacts to a more limited extent at select genes, gene transcription is less dependent on Menin in such cases. The intragenic Menin-bound enhancers observed in *ARID1B*, *CDK6* and *PAN3* (Figure 1D) also had more dramatically reduced contacts with the promoter in SEM cells (Figure S3E).

### Collapse of intragenic-bound enhancers downregulates key genes

To further characterize Menin-bound enhancers and their impact on gene regulation in SEM (MLL::AF4) cells, we focused on decreased Menin-bound contacts. The majority (61%) of these were within introns or other promoters (Figure 3A), and were also enriched for Menin binding in infant ALL patient samples with MLL::AF4, but not in OCI-AML3 cells or our adult NPM1c patient sample (Figure S4A).

**Figure 3.:**
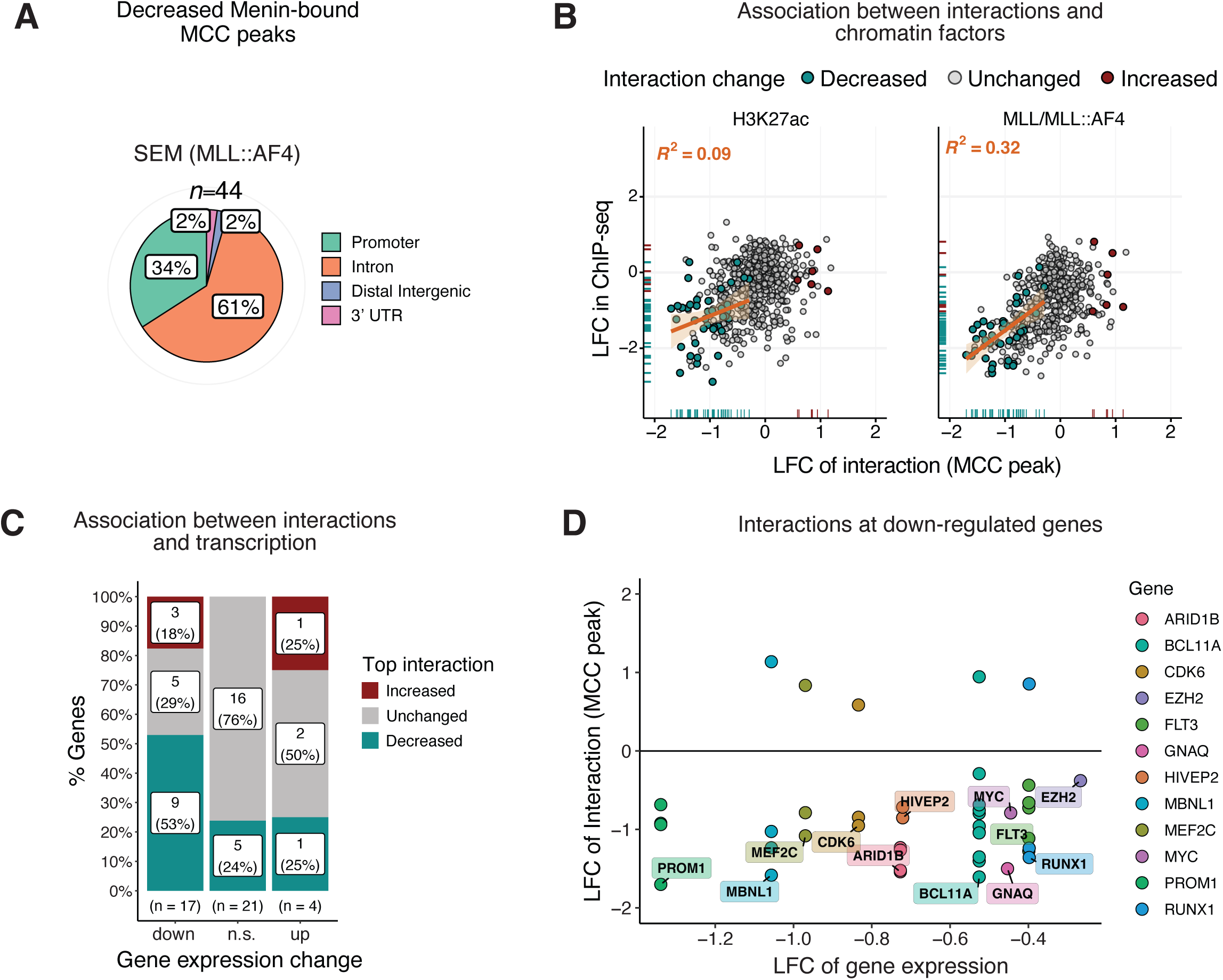
Collapse of Menin-bound enhancers is associated with decreased transcription at key genes. (A) Annotation of decreased Menin-bound enhancers in SEM cells. (B) Scatter plots showing the change in interaction frequency correlated with the change in H3K27ac or MLL/MLL::AF4 enrichment after 24 h of MENi (250 nM) in SEM cells. Colored data points are sites with a significant (MWU FDR q-value < 0.05) change in interaction. A linear fit (orange line) is shown specifically for sites of decreased interaction. (C) Association between gene expression and enhancer-promoter interaction frequency following MENi (250 nM, 24 h) in SEM cells. For each gene with multiple MCC peaks, the most statistically significant promoter interaction is presented. (D) Genes with a significantly decreased enhancer-promoter contact and decreased expression after 24 h MENi (250 nM).

Given that MENi led to both decreased H3K27ac and MLL::AF4 at MCC peaks with high levels of Menin binding in SEM cells (Figure S3B), we sought to examine the relationship between these factors and interaction frequency. At decreased interaction sites (Figure 3B, cyan data points), the loss of interaction frequency correlated more with reduced MLL::AF4 enrichment, although this correlation was still weak (R^2^ = 0.32). Prior to inhibition, decreased interaction sites had higher Menin enrichment relative to unchanged peaks (Figure S4B), suggesting that these sites may be more sensitive to Menin levels to stabilise MLL::AF4 and contacts with the promoter.

To confirm that Menin-bound enhancer-promoter contacts were functionally associated with transcription, we linked the most significant interaction change to gene expression and observed that downregulated genes were more likely to have a decreased enhancer-promoter contact in SEM cells (Figure 3C). Among the genes with a concordant decrease in enhancer contacts and expression were *ARID1B, FLT3, PROM1*, and *CDK6* (Figure 3D), at which the MLL::AF4 complex has previously been shown to sustain enhancer-promoter contacts.^36,37^ This confirmed that enhancer activation and maintenance of proximity can be an important function of Menin in MLL::AF4 leukemia.

### Menin maintains patient-unique enhancer elements in infant *MLL*-rearranged patient cells

We recently showed that enhancer activity can drive transcriptional heterogeneity in ALL patient cells that all carry the MLL::AF4 fusion.^39^ The largest proportion of enhancers that varied between patients were also found within intronic elements. While a machine learning prediction model identified the MLL::AF4 fusion and the PAF1 complex as the main predictive features of heterogenous enhancer activity. Since Menin is a key component of the MLL::AF4 complex, we hypothesized that Menin inhibition may be effective in disrupting patient-unique enhancers. To test this, we performed ChIPmentation and MCC in three primary infant ALL samples with MLL fusions proteins following Menin inhibition (Figure 4A, Figure S5A). Our samples included two patients with t(4;11) (MLL::AF4, patients 9422 and 873) and one patient with a rarer^15^ t(1:11) translocation, which results in the MLL::EPS15 fusion protein (patient 1823). Patient characteristics are provided in Supplementary Table 1.

**Figure 4.:**
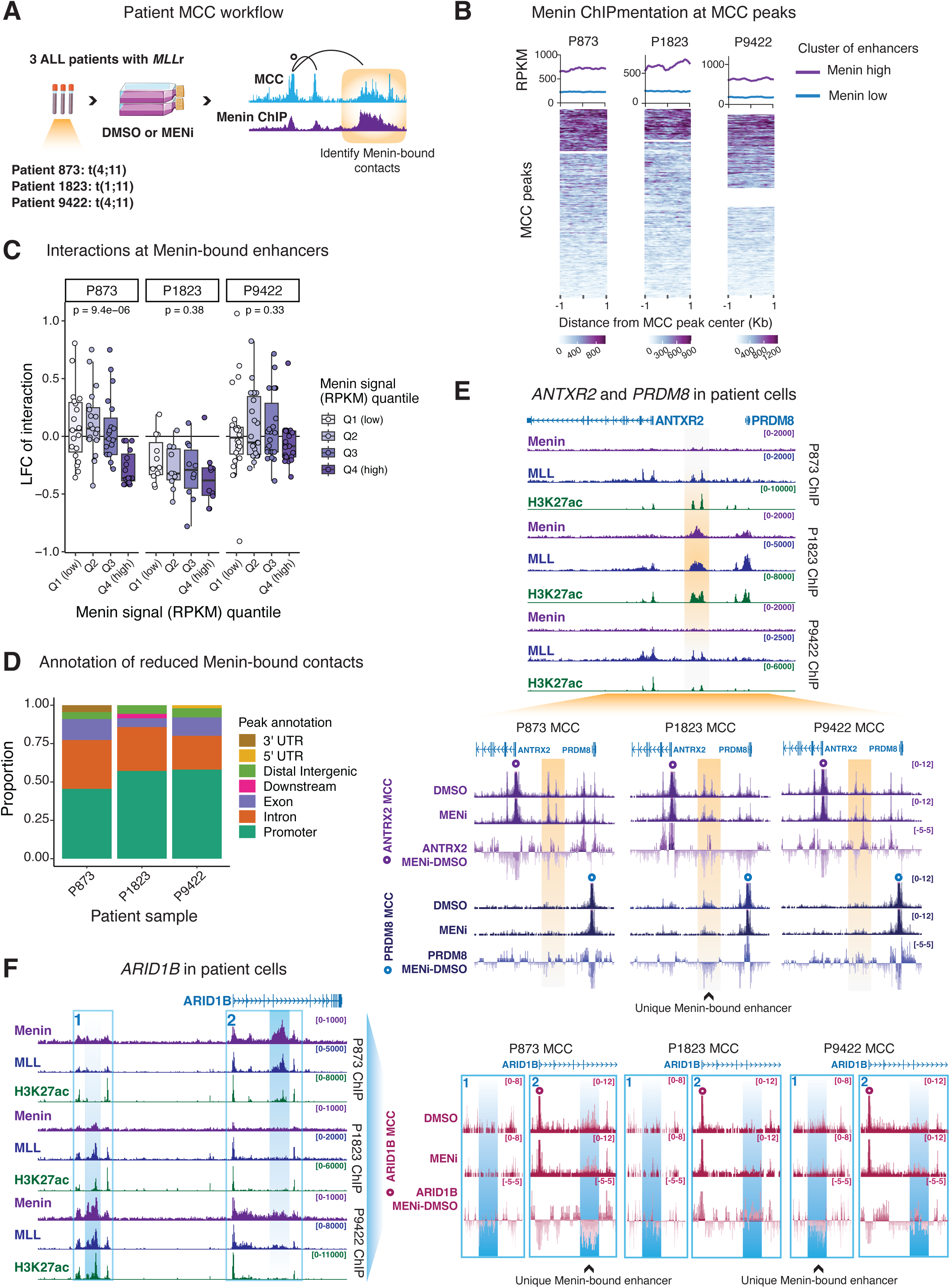
Menin-bound enhancers are sensitive to Menin inhibition in primary patient ALL cells. (A) Schematic of Micro Capture-C experiment in primary patient cells. (B) Metaplots and associated heatmaps of Menin binding at MCC peaks in patient cells, clustered by Menin binding (hclust = 2). (C) Log2 fold changes in MCC at Menin-bound enhancers stratified into quartiles by Menin signal for each patient sample. *p*-values are reported from Kruskal-Wallis tests. (E) Visualization of Menin, MLL/MLL::AF4 and H3K27ac ChIPmentation (top, mean signal) and MCC (bottom, mean+whiskers), at *ANTRX* and *PRDM8* in primary patient cells. Menin-bound sites unique to Patient 1823 are highlighted in yellow. (E) Visualization of Menin, MLL/MLL::AF4 and H3K27ac ChIPmentation (left, mean signal) and MCC (right, mean+whiskers) at *ARID1B* in primary patient cells. Menin-bound sites unique to Patient 873 and Patient 9422 are boxed in blue.

First, we confirmed that Menin was enriched at a subset of MCC peaks (Figure 4B), although this was a lower proportion than we observed in SEM cells (Figure 3B). Given heterogeneity in Menin binding and enhancer activity among the patients, we focused on comparing the effects of MENi on enhancers within each patient individually, rather than treating the patients as biological replicates and applying strict statistical thresholds (see Methods for additional details). We observed decreased (LFC < 0) interactions at some Menin-bound enhancer domains, with the largest changes in interactions tending to occur at sites with the greatest Menin enrichment at the enhancer (Figure 4C). Most of the reduced contacts mapped to other promoters (within 2 Kb of a TSS), followed by reduced enhancers within introns and exons (Figure 4D).

We observed heterogeneity in Menin binding, and consequently in enhancer usage, among the three patients. For example, Patient 1823 (P1823), who carried the MLL::EPS15 fusion, exhibited a unique Menin-bound enhancer at an intergenic site between the *ANTXR2* and *PRDM8* loci (Figure 4E, top, yellow highlight). Interestingly, the *ANTXR2* promoter contacted this region in all three patients, but the *PRDM8* promoter interacted with it exclusively in P1823. Menin inhibition specifically reduced the interaction with the *PRDM8* promoter in this the P1823 sample (Figure 4E, bottom, yellow highlights) following loss of Menin binding (Figure S5B).

The *ARID1B* locus represented another region of variation for enhancer activity and Menin binding in the patients. Although both had the MLL::AF4 fusion, P873 exhibited a uniquely large Menin-bound enhancer in intron 4 (Figure 4F, left, box 2), whereas in P9422, Menin occupied an intergenic enhancer upstream of the *ARID1B* promoter (Figure 4F, left, box 1). Following Menin inhibition, the P873 *ARID1B* promoter specifically lost interactions with the intron 4 Menin-bound site, whereas the P9422 *ARID1B* promoter lost interactions with its upstream Menin-bound enhancer, which correlated with reduced Menin occupancy (Figure S5C).

Together, these observations suggest that Menin inhibition can disrupt enhancer-promoter contacts even in heterogeneous patient cells harboring MLL::AF4 (P9422 and P873), as well as in patient cells harboring other, more rare, translocation events such as MLL::EPS15.

### Menin interactors point to its context-specific roles in transcription

We have previously shown that MLL::AF4 and interacting complexes such as DOT1L, FACT and PAF1, which are traditionally associated with transcription elongation^53–55^, can also regulate enhancer activity in SEM cells^36,37^. Consequently, we sought to interrogate whether Menin interacts with distinct protein complexes, such as additional co-activators in the presence of MLL::AF4, to facilitate distinct transcriptional processes based on the leukemia context.

To test this, we returned to the cell line models to perform endogenous Menin co-immunoprecipitation from nuclear extracts followed by mass spectrometry (IP-MS) in SEM (MLL::AF4) and OCI-AML3 (NPM1c) cells (Figure 5A). Protein abundances in the nuclear proteomes and immunoprecipitates were highly correlated (Pearson *r* ≥0.7) between SEM and OCI-AML3 cells (Figure S6A). We defined Menin interactors as proteins statistically enriched (FDR < 0.01) over IgG controls and compared this set between SEM and OCI-AML3 cells (Figure 5B). Among the most differentially enriched proteins were lineage-specific factors such as PAX5 in SEM cells and MNDA in OCI-AML3 cells. Interestingly, we also found that the acetyltransferase P300, which regulates promoter and enhancer activity via H3K27ac^56,57^, was also more enriched in the Menin interactome in SEM cells compared to OCI-AML3. As such, we hypothesised that MLL::AF4 may also co-opt Menin for driving enhancer activation in co-operation with a unique interactome of more general transcriptional co-activators.

**Figure 5.:**
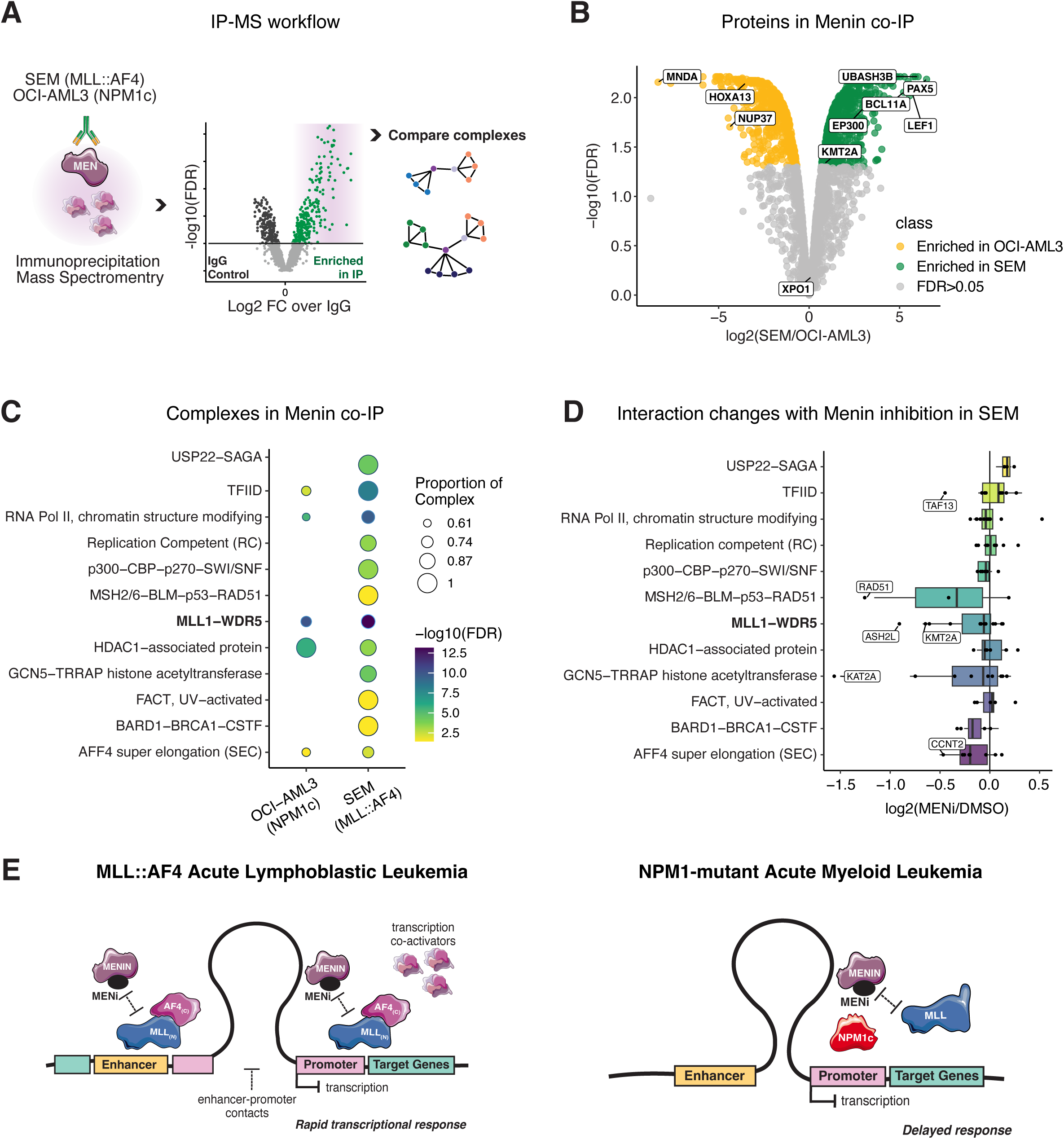
Menin associates with transcriptional regulators in leukemia cells. (A) Schematic of IP-MS workflow. (B) Volcano plot comparing statistically enriched Menin interactors in SEM and OCI-AML3 cells. Proteins were pre-filtered on enrichment over IgG controls (FDR < 0.01) in each cell line. (C) Global enrichment (FDR < 0.05) of select complexes involved in transcription in Menin co-immunoprecipitations. (D) Log2 fold changes in protein abundance in Menin co-IPs from DMSO or MENi-treated (24 h 250nm) SEM cells. (E) Model of Menin activity in MLL::AF4 and NPM1-mutant leukemia: Menin associates more strongly with transcription co-activators and stabilizes enhancer-promoter proximity to activate genes in MLL::AF4 but not NPM1c AML.

To identify general transcription complexes associated with Menin, we performed an enrichment analysis for protein complexes in the Corum database^58^ for each cell line. As expected, we found the MLL1 complex enriched in both SEM and OCI-AML3 cells (Figure 5C). However, members of FACT, SAGA, and MSH2/6 DNA repair machinery were specifically enriched in SEM cells. The expression of these complex components was highly correlated in the nuclei of both SEM and OCI-AML3 cells (Figure S6B), indicating that the observed differential enrichment reflects interactions with Menin rather than simply differential expression between these cell lines.

To assess whether the transcription-associated complexes uniquely enriched in SEM cells are also key for cell survival, we referenced DepMap gene effect scores for the genes making up these complexes. Gene effect scores, where a score of -1 represents the median of common essential genes^59^, were highly correlated in SEM and OCI-AML3 cells, with a subset emerging as essential in both cell lines (Figure S6C). The selective enrichment of these transcriptional activators in the SEM Menin interactome may explain the rapid transcriptional changes and associated short-term sensitivity to Menin inhibition, as Menin may be repurposed to stabilise the co-operation of these complexes with the MLL::AF4 complex. Menin inhibition for 24 h indeed led to decreased (LFC < 0) enrichment of some transcriptional complex components in Menin immunoprecipitations (Figure 5D), suggesting that additional factors may rely on Menin binding to chromatin to maintain their association with the MLL::AF4 complex.

### Distinct roles for Menin in MLL::AF4 and NPM1-mutant leukemia

Together, our data point to distinct Menin activity in MLL::AF4 and NPM1c acute leukemias in transcriptional activation and modulating enhancer activity (Figure 5E). In MLL::AF4 cells, Menin functions as a transcriptional co-activator, facilitating promoter–enhancer contacts and interacting with additional transcription activators. Menin inhibition therefore results in rapid, widespread changes in transcription in MLL::AF4 cells. In contrast, in NPM1c AML cells, Menin loss has a restricted acute effect on transcription at small subset of target loci, likely due to a more indirect effect on stabilizing NPM1c binding rather than directly interacting with transcription elongation complexes. These differences in Menin protein complex composition and Menin activity at enhancers could explain the greater transcriptional disruption observed with MENi in MLL::AF4 compared to NPM1c cells.

## Discussion

Menin inhibition is a promising targeted therapy for high-risk *MLL*-rearranged and *NPM1*-mutant acute leukemias, which both depend on Menin for the expression of key genes and cell survival. However, our data highlight that the mechanism by which Menin affects aberrant transcription differs between the two diseases. Upon MENi, SEM cells (MLL::AF4 ALL) exhibited rapid transcriptional dysregulation, which affected thousands of genes, and loss of intragenic Menin-bound enhancer activity. Menin-bound enhancers were similarly affected in primary patient cells with MLL::AF4 and MLL::EPS15, despite enhancer heterogeneity. On the other hand, OCI-AML3 cells (NPM1c AML) showed transcription changes in approximately one hundred genes at the same timepoint, while enhancer activity was unaffected. Therefore, the major contribution of Menin to transcription is highly context-dependent, and linked to its enhancer activity and the protein complexes it participates in.

Recently, we found that leukemia samples from individual patients often have unique enhancer landscapes, and that these unique enhancers are sometimes linked to important oncogenes that drive disease aggressiveness.^39^ Some *MLL*r patients fail to respond to Menin compounds in clinical trials, so one possibility is that they have unique enhancers able to maintain oncogene expression despite disruption of Menin activity. However, here we show that even patient-specific enhancers are sensitive to Menin disruption, losing contacts with their target promoters. This reveals a fundamental dependency of enhancer activity on Menin’s role within the MLL fusion protein complex, demonstrating that these unique enhancers cannot operate independently of Menin. Taken together, these findings point to a critical insight: epigenetic mechanisms of Menin resistance are far more likely to emerge through alterations in the function of the complex itself, rather than through the formation of new regulatory enhancers.

Our Menin co-immunoprecipitations from SEM and OCI-AML3 cells revealed that Menin interacts with transcription regulators with some cell-type specificity. This adds to a complex body of literature illuminating context-specific Menin interactions that can trigger either transcription activation or repression^9^. Focusing on transcription regulators, we found that P300 and the FACT complex were more enriched in the Menin interactome of SEM cells compared to OCI-AML3 cells. Given that loss of either PAF1, FACT ^36^ or H3K79me2/3^37^ can lead to decreased enhancer promoter interactions at MLL::AF4-bound enhancers, these multiple vulnerabilities suggest that the overall stability of the MLL::AF4 complex, rather than one specific component, may be key to the maintenance of enhancer-promoter contacts and transcription. This points to a possible role for Menin in stabilizing these factors at enhancers in MLL::AF4 ALL cells. Understanding how these complexes form in the presence and absence of Menin binding could be key to targeting the emergence of epigenetic resistance mechanisms in patients.

Both MLL-FPs and NPM1c are transcriptional activators that require Menin binding at a subset of their target promoters. Wang *et. al.* (2023) showed that chromatin-associated NPM1c forms transcriptional hubs, which include elongation factor ENL, Menin, and RNA Polymerase II, through multivalent interactions^24^. However, they also showed that NPM1c did not alter chromatin or trigger new transcription when introduced de novo; rather it further upregulates genes with an established baseline expression. MLL-FPs on the other hand, in addition to maintaining expression of active genes via the recruitment of co-activators and elongation factors^45,46^, can also activate self-renewal genes characteristic of hematopoietic stem cells when exogenously expressed^60^. Recently, it was demonstrated that MLL-AF10 can activate new enhancer elements^40^. Therefore, destabilizing MLL::AF4, a more potent and wide-spread transcriptional activator compared to NPM1c, via MENi could be more disruptive to the oncogenic gene expression programme, indicating that that Menin is co-opted as a transcriptional activator to a greater extent than when in a complex with NPM1c.

While our analysis focused on MLL::AF4 ALL, it has also previously been observed that leukemia cell identity can dictate the response to MENi. Comparing 4-day treatments with Revumenib, a Menin inhibitor used in the clinic, in *MLL*r AML and ALL cells has revealed that MENi rapidly induces apoptosis and cell cycle arrest in *MLL*r-ALL cells, whereas in *MLL*r-AML cells the response is slower and involves differentiation rather than cell death^61^. Therefore, understanding Menin activity in the presence of both different driver mutations and in different leukemia subtypes is key to extending MENi as a therapeutic option to the right patient populations^62^. Following evidence that Menin can regulate the transcriptional activity of NUP98 fusion proteins^63^ and oncoproteins arising from *UBTF* tandem duplications^64^, patients with these lesions are now included in Menin inhibitor clinical trials (e.g. NCT05326516, NCT05360160, NCT04811560, NCT06177067, APAL2020K ITCC-101/COG/PEDAL).

Unexpectedly, Menin and its interacting complexes are also an emerging vulnerability in solid tumors^65,66^. How Menin regulates genes in cancers with other driver mutations, and whether its function goes beyond supporting the binding of other oncoproteins on chromatin, remains to be elucidated.

Monitoring the transcriptional responses to MENi over time could also be key to classifying patient responses as we ascertain resistance mechanisms. The AUGMENT-101 clinical trial reported that mutations in the Revumenib binding pocket on Menin can restore Menin/MLL-FP binding and expression of MLL-FP target genes^67^. However, when resistance arose without new mutations, the leukemias persisted despite the expected downregulation of Menin targets. This suggests that some leukemias can cease to depend on the Menin/MLL-FP transcriptional program despite an initial and sustained response to MENi. Distinct transcriptional responses to MENi, and how they link to clinical outcomes, can therefore provide a rationale for subtype-specific combination therapies.

## Acknowledgements

We thank Jerry McGeehan (Syndax Pharmaceuticals) for supplying the VTP50469 used in these experiments. V.S. was funded by the Wellcome Trust. A.D. and B.M.K. were supported by the Chinese Academy of Medical Sciences (CAMS) Innovation Fund for Medical Sciences (CIFMS), China (grant number: 2024-I2M-2-001-1). T.A.M. was supported by Medical Research Council (MRC, UK) Molecular Haematology Unit grant MC_UU_00029/6. The work was also supported by Blood Cancer UK grant 24016. N.T.C. was supported by a Kay Kendall Leukemia Fund Intermediate Fellowship (KKL1443). A.R. is supported by a Wellcome Trust Clinical Research Career Development Fellowship (216632/Z/19/Z) and Medical Research Council (MRC, UK) Molecular Haematology Unit grant MC_UU_00029/7. Samples and data used in this study were provided by VIVO Biobank, supported by Cancer Research UK & Blood Cancer UK (Grant no. CRCPSC-Dec21\100003). Scientific illustrations were adapted from Bioicons.com under creative commons licence CC-BY.

## Author Contributions

T.A.M, N.T.C., B.M.K., and V.S. conceived the experimental design. V.S., A.D., S.S.H., and A.L.S. carried out experiments. V.S, C.C., F.H.L., and J.C.H. analyzed and curated the data. V.S., C.C, N.T.C., and T.A.M. interpreted the data and wrote the manuscript. N.D. contributed toward MCC experimental design. I.V and S.S.H. developed mass spectrometry methods. R.W.S., M.K., and A.R. provided primary patient material. B.M.K. and T.A.M., provided funding and supervision. All authors reviewed the manuscript.

## Data Availability

Genomics data have been deposited in the Gene Expression Omnibus (GEO) as a series under the accession number GSE312345. Mass spectrometry data have been deposited to the ProteomeXchange Consortium via the PRIDE partner repository with the dataset identifier PXD071336. Publicly available cell line data and previously deposited patient data are available in GEO under accession codes: GSM5114291; GSM5139719; GSM5139645; GSM5139689; GSM8657204; GSM8657208; GSM8657184; GSM8657191; GSM5858383; GSM3635683.

## Methods

### Cell culture

SEM cells (DSMZ) were grown in IMDM supplemented with 10% FBS and 1X GlutaMax (Gibco). RCH-ACV and OCI-AML3 cells were grown in RPMI supplemented with 10% FBS and 1X GlutaMax (Gibco). Patient cells were cultured in RPMI supplemented with 5 mg/ml insulin, 5mg/ml transferrin, 5 ng/ml sodium selenite (ITS, Gibco), 200 µg/ml gentamycin, and 20% FBS.

### Patient samples

nfant (aged <1 year at diagnosis) ALL samples iALL28349 and iALL863388 were obtained from Blood Cancer UK Childhood Leukaemia Cell Bank (now VIVO Biobank, United Kingdom) under their ethics approval (REC: 23/EM/0130) and from Our Lady’s Children’s Hospital, Crumlin, Ireland (REC: 21/LO/0195). The adult AML sample was obtained from MD Anderson Centre, USA, under RB protocol LAB-01-473. Primary *MLL-*rearranged infant ALL patient specimens P1823, P873 and P9422 were collected as part of the INTERFANT studies;^68^ patient characteristics are listed in Supplementary Table 1. Informed consent was obtained from parents or legal guardians to use excess diagnostic material for research purposes, as approved by the institutional review board. These studies were conducted in accordance with the Declaration of Helsinki. All samples were processed and cultured as previously described^69^. Samples were obtained at diagnosis and contained >90% blasts.

### CellTiter-Glo

Cells were treated with a dilution series of VTP50469 fumarate (Syndax Pharmaceuticals) in triplicate wells in an opaque 96 well plate. After 48 h, CellTiter-Glo 2.0 reagent (Promega G9241) was added to each well in equal volume to the media. The plate was shaken for 2 min and incubated for 5 min at room temperature, after which luminescence was recorded on a BMG FLUOstar OPTIMA plate reader. The luminescence signal was normalised to DMSO-treated cells and averaged across replicates. The dose response curves were fitted using the R package drc^70^.

### Colony Assay

In duplicate, cells were treated with either 50 nM or 250 nM VTP50469, or DMSO, for 90 min, then added to MethoCult (StemCell Technologies H4100) containing 50 nM or 250 nM VTP50469 or DMSO. Methocult cultures were aliquoted into three replicate 3 cm dishes, for a total of 5 replicate plates per condition. After 10-14 days, the number of colonies was counted under a light microscope and the counts across replicates were averaged.

### Chromatin Immunoprecipitation followed by Sequencing (ChIP-seq)

For Menin, NPM1c and MLL-N ChIP, cells were fixed for 30 min with 2 mM Di(N-succinimidyl) glutarate (DSG) followed by 30 minutes with 1% formaldehyde. For histone modifications, cells were fixed for 10 minutes with 1% formaldehyde. Fixed cells were lysed in SDS lysis buffer (1% SDS, 10 mM EDTA, 50 mM Tris-HCl, pH 8 with protease inhibitor cocktail) and the lysate was syringe-passaged through a 27-gauge needle. The cell lysates were sonicated using a Covaris ME220 (peak power 75, duty factor 25 %, 500 cycles per burst) for 10 min for Menin, NPM1c and MLL-N ChIP, or for 5 min for histone ChIP. The sonicated lysates were pre-cleared with a 1:1 mix of Protein A:Protein G beads for 15 minutes at 4°C, then incubated with antibody overnight. Antibodies are provided in Supplementary Table 2. Five percent of the lysate was set aside as input. The following day, a 1:1 mix of Protein A:Protein G beads was added to each IP and samples were rotated at 4 °C for 3 hours. The beads were then washed three times with RIPA buffer (50 mM HEPES-KOH, pH 7.6, 500 mM LiCl, 1 mM EDTA, 1% NP-40, 0.7% Na-deoxycholate) and once with TE/50 mM NaCl. The chromatin was eluted from the beads with SDS lysis buffer for 30 min at 65 °C, then treated with RNase A for 30 min at 37 °C, followed by Proteinase K treatment combined with cross-link reversal overnight at 65 °C. DNA was purified using the ChIP DNA Clean & Concentrator kit (Zymo 5201). Libraries for sequencing were prepared using the NEBNext Ultra II library prep kit (E7645) and sequenced by 40 bp paired-end sequencing with a NextSeq500 (Illumina).

### ATAC-seq

ATAC-seq was performed as described by Buenrostro *et al*^71^. 50,000 cells were lysed in 50 µL cold lysis buffer (10 mM Tris-HCl pH 7.4, 10 mM NaCl, 3 mM MgCl2, 0.1 % IGEPAL CA-630) and centrifuged at 500 g for 10 minutes at 4°C. The pellet was resuspended in transposase buffer (1X Illumina Tagment DNA Buffer - 20034197) with 2.5 µL Tn5 Transposase enzyme and incubated for 30 min at 37°C. Transposed DNA was extracted using the MinElute PCR Purification Kit (Qiagen 28004) following the manufacturer’s instructions, then PCR-amplified using Nexterra indices and NebNext Ultra II Q5 MasterMix (NEB M0544S) for a total of 12 cycles. Amplified tagmented DNA was purified with the MinElute PCR Purification Kit and sequenced by 75 bp paired-end sequencing.

### Transient Transcriptome RNA-sequencing (TT-seq)

TT-seq was performed as previously described^72^. In triplicate, 4-5x10^7^ cells were labelled with 500 µM 4-thiouridine (Sigma) for 5 min. 50 ng spike-in RNA, consisting of three 4-thiouridine-labelled transcripts from a CAS9-encoding vector, was added to each replicate. Total RNA was extracted with either Trizol or RNeasy Maxi kit (Qiagen), DNase I-treated, and fragmented by sonication on Covaris ME220 for 10 s (peak power 70 W, duty factor 20%, 1000 cycles/burst). 150-200 µg input RNA was labelled with EZ-Link HPDP-Biotin (Pierce 21341) for 90 min and purified by chloroform/isoamylalcohol extraction. Labelled RNA (i.e. nascent transcripts) was isolated using the uMACS Streptavidin Kit (Miltenyi 130-074-101) following the manufacturer’s instructions and purified with the RNeasy MinElute Cleanup Kit (Qiagen 74204). cDNA synthesis and library preparation for sequencing were performed using the NEBNext Ultra II Directional RNA Library Prep Kit (NEB E7760) following the protocol for purified mRNA. The libraries were quantified using the KAPA Library Quantification Kit (Roche KK4824) and sequenced by 80 bp paired-end sequencing or 150 bp paired-end sequencing with a NextSeq500 (Illumina).

### ChIPmentation

ChIPmentation was performed as previously described^73^. Protein A magnetic beads were conjugated with 1.5 µg of antibody for 4 hours at 4 °C with rotation in 150 µL PBS 0.5% BSA 1 x protease inhibitor cocktail (PIC). Antibodies are provided in **Supplementary Table 1.** Cells were fixed as for ChIP-seq. Fixed cells were lysed in 120 µL Lysis Buffer (50 mM Tris-HCl pH 8.0, 0.5% SDS, 10 mM EDTA, 1x protease inhibitor cocktail (PIC) and sonicated using Covaris ME220 using the same conditions as for ChIP-seq to generate 200-300 bp fragments. Triton-X100 was added to the sonicated chromatin to a final concentration of 1% and incubated for 10 min at room temperature to neutralize the lysis buffer SDS. The chromatin was pre-cleared with protein A beads for 30 minutes at 4 °C with rotation. Antibody-conjugated beads were washed with 150 µL PBS 0.5% FBS, then added to the pre-cleared chromatin and incubated overnight at 4 °C with rotation. The following day the beads were washed three times with RIPA buffer (as in ChIP-seq), once with Tris-EDTA and once with 10 mM Tris-HCl pH 8.0. The beads were then resuspended in Tagmentation Buffer (10 mM Tris-HCl pH 8.0, 5 mM MgCl_2_, 10% dimethylformamide) with 1 µL Tn5 transposase (Illumina). Samples were incubated at 37°C for 5 minutes, after which the transposition reaction was stopped with 150 µL RIPA buffer. Beads were washed with 10 mM Tris-HCl pH 8.0. PCR was performed with NEBNext Ultra II Q5 Mastermix and Nextera index primers (125 nM final concentration) for 13 cycles. AMPure XP beads (Beckman A63881) in a 1:1 ratio were used for clean-up. Samples were sequenced with paired-end sequencing with a NextSeq500 (Illumina).

### Immunoprecipitation

4x10^7^ cells were lysed in 1 mL Buffer A (10 mM HEPES pH 7.9, 10 mM KCl, 1.5 mM MgCl_2_, 0.34 M sucrose, 10% glycerol) with 0.2% NP-40 and 1x PIC for 5 minutes on ice. Following a 5 minute centrifugation at 500g and 4 °C, the cytoplasmic fraction (supernatant) was removed and intact nuclei were washed once in Buffer A + PIC. The nuclei were resuspended in Nuclear Lysis Buffer (50 mM Tris-HCl, pH 8, 2 mM MgCl_2_, 150 mM NaCl, 0.5% NP-40, 1x PIC) with 250 U benzonase (Sigma) and rotated for 1 h at 4 °C. For cells treated with 250 nM MENi for 24 h, all buffers were supplemented with VTP50469 to 1mM. After centrifugation at 20,000 g for 10 min and 4 °C, 2% of the supernatant was set aside as a nuclear input and the remainder was used for immunoprecipitation with 4 µg anti-Menin antibody (Bethyl A300-105A) overnight at 4 °C. An equal amount of rabbit IgG antibody (Cell Signalling DA1E) was used in parallel as a control. A 1:1 mix of Protein A:Protein G beads was added to each IP and rotated at 4 °C for 3 h. The beads were washed three times with Nuclear Lysis Buffer and once with TE/50 mM NaCl. The beads were then resuspended in 35 µL 2X LDS buffer and proteins were eluted by incubating at 95 °C for 5 min. For quality checks, approximately 4% of the sample was used for SDS-PAGE followed by Silver Stain (Pierce^TM^ Silver Stain Kit, ThermoFisher 24612) and 8.5% was used for western blotting for LEDGF, a protein known to interact with Menin.

### Mass Spectrometry

Peptides were prepared for Mass Spectrometry with the S-trap micro spin column digestion protocol (PROTIFI), following the manufacturer’s instructions. Briefly, samples were reduced, alkylated and acidified, then purified with the S-Trap micro column. After 5 washes with the binding/wash buffer, 5 µg Trypsin was added and columns were incubated at 37 °C overnight. Peptides were resuspended in 5% formic acid and 5% DMSO and then trapped on an Acclaim™ PepMap™ 100 C18 HPLC Columns (PepMapC18; 300µm x 5mm, 5µm particle size, Thermo Fischer) using solvent A (0.1% Formic Acid in water) at a pressure of 60 bar and separated on an Ultimate 3000 UHPLC system (Thermo Fischer Scientific) coupled to a QExactive mass spectrometer (Thermo Fischer Scientific). The peptides were separated on an Easy Spray PepMap RSLC column (75µm i.d. x 2µm x 50mm, 100 Å, Thermo Fisher) and then electro-sprayed directly into an QExactive mass spectrometer (Thermo Fischer Scientific) through an EASY-Spray nano-electrospray ion source (Thermo Fischer Scientific) using a linear gradient (length: 60 minutes, 5% to 35% solvent B (0.1% formic acid in acetonitrile and 5% dimethyl sulfoxide), flow rate: 250 nL/min). The raw data was acquired in the mass spectrometer in a data-independent mode (DIA). Full scan MS spectra were acquired in the Orbitrap (inclusion list with scan range 495 to 995 m/z, 20m/z increments, with an overlap of +/- 2 Daltons, resolution 35000, AGC target 3e6, maximum injection time 55ms). After the MS scans peaks were selected for HCD fragmentation at 28% of normalised collision energy(NCE) / stepped NCE. HCD spectra were also acquired in the Orbitrap (resolution 17500, AGC target 1e6, isolation window 20m/z).

### Micro Capture-C

Micro Capture-C was conducted as previously^52,74^. In biological triplicates, 5x10^7^ SEM and OCI-AML3 cells were treated with 250 nM VTP50469 or DMSO for 24 hours. Each patient sample was also divided into DMSO or 250 nM VTP50469 for 24 hours, for one biological replicate per treatment condition. Cells were fixed with 2% formaldehyde for 10 min, followed by quenching with glycine and a PBS wash. Cells were then aliquoted into three technical replicates. 0.0005% digitonin (Sigma) was used to permeabilize the cells for 15 minutes, after which they were snap frozen in three aliquots. After thawing, 16 x 10^6^ cells (one aliquot) were resuspended in 10mM Tris-HCl pH 7.5, 1 mM CaCl_2_ and divided into three micrococcal nuclease (NEB) digestion reactions with 7.5, 10 or 12.5 Kunitz U for 1 h at 37 °C shaking at 550 rpm in a Thermomixer (Eppendorf). MNase digestion was stopped with 5 mM EGTA and an aliquot was set aside to assess digestion by TapeStation. The cells were washed with PBS/5 mM EGTA and resuspended in: DNA ligase buffer, 400 µM dNTPs, 5 mM EDTA, 100 U/µL DNA Polymerase 1, Large (Klenow) Fragment (NEB), 200 U/µL T4 PNK (NEB) and 300 U/µL T4 DNA ligase (Thermo Scientific). This ligation reaction was incubated at 550 rpm at 37°C for 2 h followed by 20 °C for 8 hours, then held at 4 °C in a ThermoMixer. Cells were then washed with PBS and digested with Proteinase K at 65 °C overnight to simultaneously reverse crosslinks. DNA was purified using phenol:chloroform:isoamyl alcohol (25:24:1) and digestion and ligation were assessed by TapeStation (Agilent D1000). For SEM and OCI-AML4 cells, two 3C libraries (technical replicates) per biological replicate were sonicated to 200 bp on Covaris ME220. For patient samples, four 3C libraries (technical replicates) per treatment condition were sonicated. Indexing was performed in duplicate using Illumina paired-end adaptors, NEB Indices and Herculase II (Agilent). Enrichment for promoters of interest was performed twice with 120 nt biotinylated capture probes using the HyperCapture Target Enrichment Kit (Roche), using M270 streptavidin beads (Invitrogen) for the pulldown. Capture probes were designed using the Capsequm pipeline (https://github.com/Davies-Genomics-and-Genome-Editing-Lab/oligo_design_pipeline). Sequences are provided in Supplementary File 1. Samples were sequenced by 150 bp paired-end sequencing on NextSeq500 (Illumina).

### Bioinformatic analysis

Publicly available genomics data was accessed from GEO. References to the data are provided in Supplementary Table 3.

#### ChIP-seq and ChIPmentation

Sequencing reads were trimmed using trim_galore (https://github.com/FelixKrueger/TrimGalore) and aligned to hg38 using bowtie2.^75^ DeepTools^76^ was used to create bigwig files for visualization on UCSC and to generate heatmaps and metaplots. HOMER^77^ was used to call peaks using findPeaks and the flag - style histone. ChIP-seq signal at peaks was quantified using the deepTools command multiBigwigSummary BED-file. Differential peak analysis was performed with the R package DiffBind.^78^ Heatmaps of differential peaks were generated with R package ComplexHeatmap.^79^

#### TT-seq

Transient transcriptome RNA sequencing reads were aligned to human genome (hg38) using STAR,^80^ and gene expression levels were determined using the Subread package tool featureCounts. ^81^ Differential gene expression analysis was completed using edgeR.^82^ Metaplot visualisations of strand-specific transcription were created using the deepTools command computeMatrix and plotted in R using ggplot2. χ^2^ tests were performed using the R package ggstatsplot.^83^

#### Genomic annotation

ATAC, ChIP-seq and ChIPmentation peaks were annotated with the R package ChIPseeker.^84^ Promoters were defined as ATAC-seq peaks that overlap H3K27ac peaks and are within 2 kb of an annotated transcription start site (TSS). Enhancers were defined as ATAC-seq peaks that overlap H3K27ac peaks and are 2 kb or more away from an annotated TSS. In heatmaps and metaplots depicting enhancers, ChIP-seq signal was centred on the enhancer ATAC peak for cell lines. For patient samples missing ATAC data, the ChIP-seq signal was centred on H3K27ac peaks. Enhancers were assigned to genes using the nearest annotated TSS.

#### Immunoprecipitation Mass Spectrometry

Data search was performed in a library-free search in DIA-NN^85^ 1.8.1 with the database UPR UPR_Homo sapiens_9606_UP000005640_20200803.fasta. FDR (false discovery rate) was set to 0.01 and MBR (match between runs) was enabled. Mass spec intensities were normalized by MaxLFQ (generic label free technology) and transformed to log2. Imputation was performed in R by replacing missing values with randomly sampled values from a down-shifted normal distribution following methods described in ^86^. Keratins and common contaminants identified in 95% of the CRAPome database^87^ were excluded from the data. Comparisons between conditions (IgG and Menin IPs) were performed with moderated t-tests using the R package Genoppi^88^ (https://github.com/lagelab/Genoppi) and corrected for multiple comparisons using False Discovery Rate with the R package fdrtools.^89^ Volcano plots were generated using ggplot2. Enrichment analyses were performed with a hypergeometric test from the Genoppi package using the CORUM database.^58^

#### Micro Capture-C

Analysis was performed using the Micro Capture-C pipeline from the Davies lab (https://github.com/jojdavies/Micro_Capture-C). Peak calling was performed with deep-learning-based peak caller LanceOtron.^90^ The peaks were filtered using a custom script from the Davies Lab based on size, width, and distance from the capture viewpoint. Unique junctions were counted at each MCC peak, then normalized using cis unique ligation junctions for the corresponding viewpoint as previously reported.^52^ For the analysis of cell lines, junctions were compared between conditions using a Mann-Whitney U test corrected for multiple comparisons using R package fdrtools, and comparisons were visualized in R using ggplot2. MCC peaks were intersected with Menin ChIP-seq peaks using bedtools intersect. Pearson’s chi-square test was used to test for independence between Menin peaks and differential MCC peaks.

For MCC in patient samples (Figure 4), MCC peaks were intersected with Homer Menin peaks that were within 2 Mb of the capture viewpoint (± 1 Mb each side), excluding a ± 2 kb region around the center of the viewpoint. LaneOtron MCC peaks often failed to capture broad Menin-bound enhancers; as a result, unique ligation junctions were counted over Menin peaks that intersected with MCC peaks, specific to each patient. Each patient had a unique set of MCC (4 technical replicates per condition) and Menin peaks (1 replicate per condition), and was treated independently. Log2 fold changes in ligation junction counts were computed per peaks for each patient using the average counts from 4 technical replicates.

#### Genomic track visualization

BigWigs were visualized in the UCSC Genome Browser (genome.ucsc.edu). The signal visualized is the mean or mean + whiskers (1 standard deviation, and maximum/minimum) where indicated. Subtraction plots are smoothed to 10 pixels.

### Use of Artificial Intelligence

ChatGPT Edu was used for troubleshooting and optimizing bash and R code generated by the authors. AI was not used in writing the manuscript.

---

**Supplementary Figure 1.:**
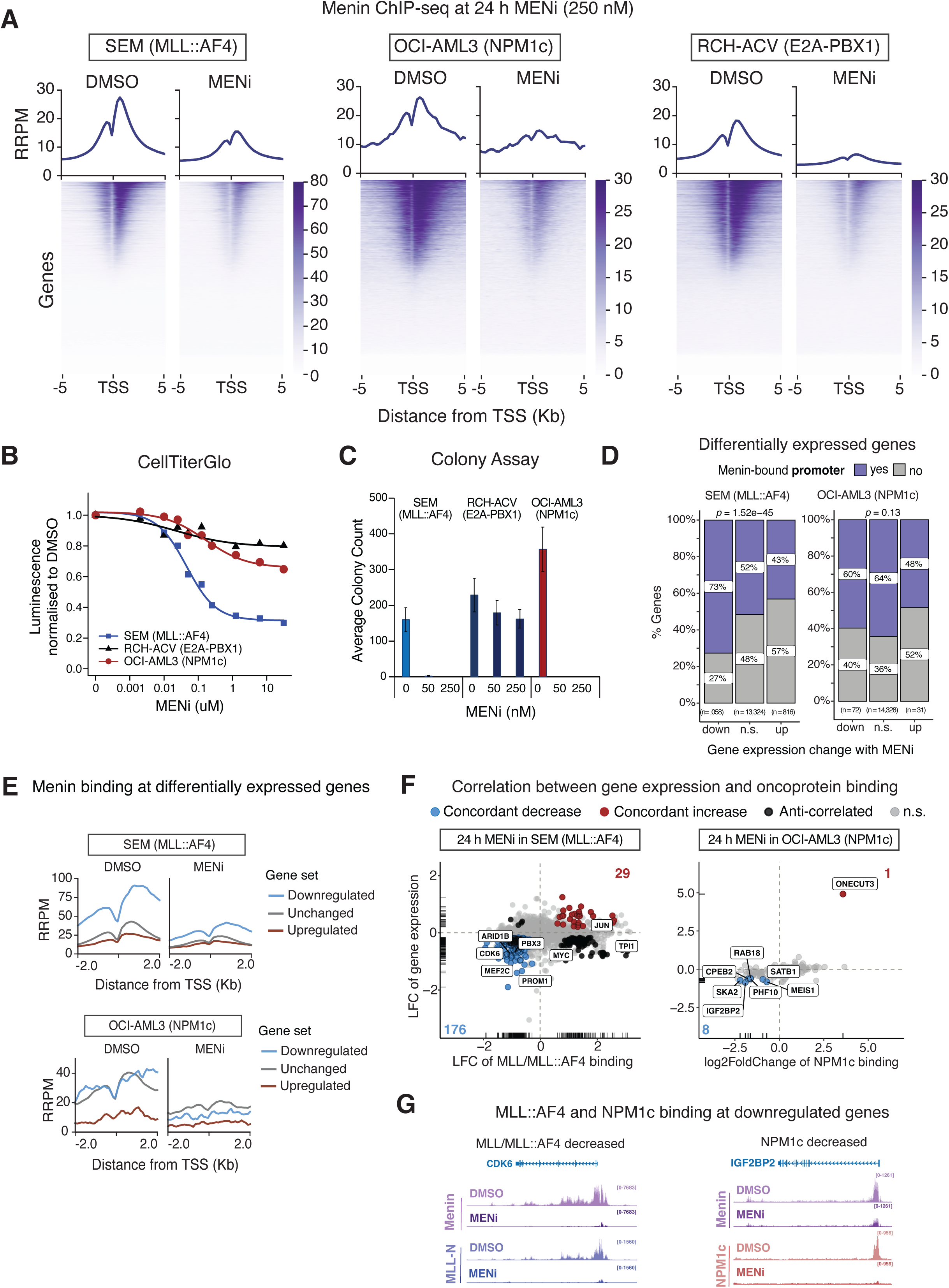
Association between Menin binding, cell viability, and transcription changes. (A) Menin ChIP-seq after 24 h treatment with DMSO or 250 nM Menin inhibitor VTP50469 (MENi) in SEM, OCI-AML3, and RCH-ACV cells. Metaplots show mean reference normalized reads per million (RRPM) centered on transcription start sites (TSS) genome-wide. (B) Dose response curves to MENi recorded at 48 h using CellTiterGlo (n = 5 for SEM and RCH-ACV cells, n = 3 for OCI-AML3 cells). (C) Colony forming units recorded after culture in Menin inhibitor (n=5). Error bars represent standard error. (D) Proportion of differentially expressed genes (n.s. = not significant) following MENi with Menin binding at the promoter (± 2 kb around the TSS). Reported *p*-values are from a Pearson chi-square test. (E) Mean Menin binding (RRPM = reference normalised ChIP-seq reads per million) in SEM and OCI-AML3 cells at promoters of differentially expressed genes in DMSO or MENi-treated cells. (F) Scatter plots correlating changes in gene expression and changes in MLL/MLL::AF4 or NPM1c binding following 24 h MENi (250nM). Genes are colored if they display a significant alteration (p-adj < 0.05) in both expression and MLL::AF4/NPM1c binding. (G) Examples of altered Menin, MLL/MLL::AF4, and NPM1c binding following 24 h MENi (250nM) in SEM cells (left) and OCI-AML3 cells (right).

**Supplementary Figure 2.:**
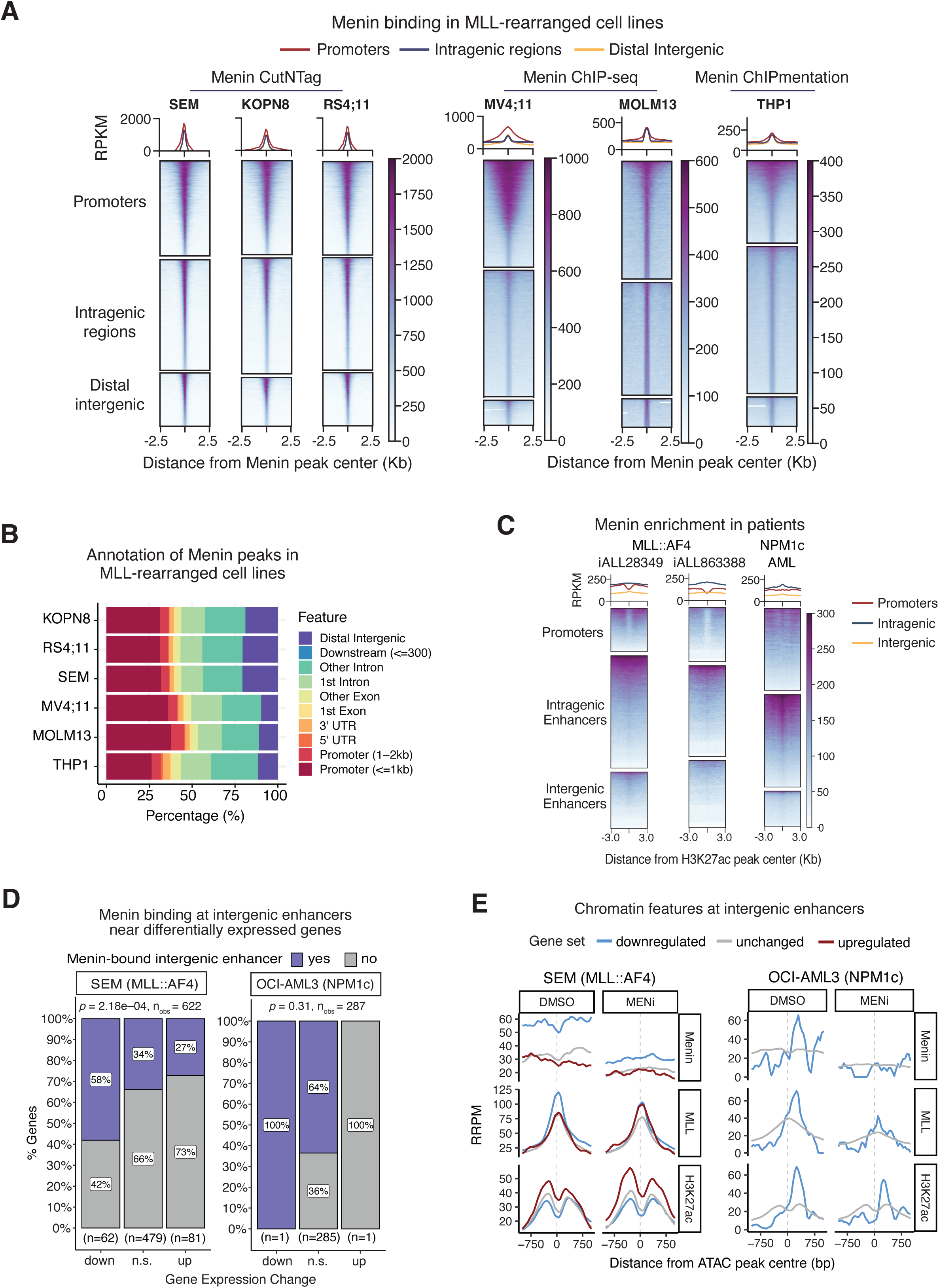
Menin binds enhancer elements. (A) Heatmaps and metaplots of mean Menin signal at promoters (±2 kb of a TSS), intragenic regions, and distal intergenic regions in cell lines with *MLL* translocations. (B) Annotation of Menin peaks in cell lines with *MLL* translocations. (C) Heatmaps of Menin ChIPmentation at promoters and enhancers, centred on H3K27ac peaks, in primary samples from two infant MLL::AF4 ALL patients (iALL28349, iALL863388) and one adult NPM1-mutated AML patient. (D) Association between Menin binding at intergenic enhancers and gene expression following 24 h of MENi (250 nM). *p*-values are reported from a Pearson chi-squared test. (E) Metaplots representing mean Menin, MLL, and H3K27ac signal at intergenic enhancers annotated to differentially expressed genes after 24 h of MENi (250 nM) in SEM and OCI-AML3 cells.

**Supplementary Figure 3.:**
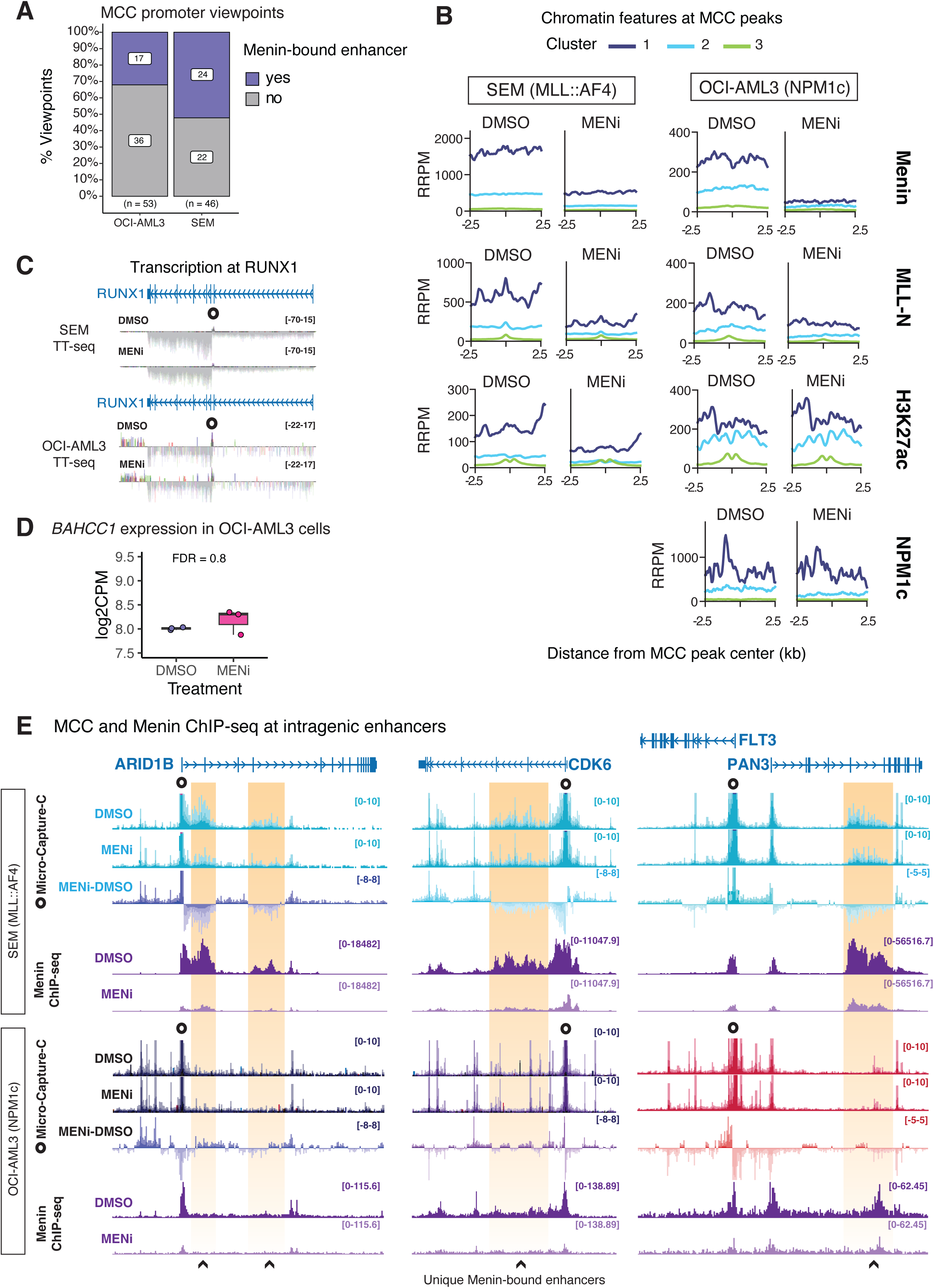
Micro-Capture-C reveals Menin-sensitive enhancers in SEM cells. (A) Proportion of genes selected for Micro-Capture-C promoter viewpoints annotated with whether they have a proximal Menin-bound enhancer, assigned by nearest TSS. (B) Mean Menin, MLL/MLL::AF4, H3K27ac, and NPM1c signal at MCC peaks in SEM cells and OCI-AML3 cells, clustered with k-means = 3 across all ChIP-seq samples per cell line. (C) Visualisation of TT-seq data at RUNX1 in SEM and OCI-AML3 cells. The circle denotes the MCC capture viewpoint. (D) *BAHCC1* expression after DMSO or 24 h MENi (250 nM) in OCI-AML3 cells, TT-seq data (CPM = counts per million). (E) Visualization of MCC (mean with whiskers) and Menin ChIP-seq (mean) data at *ARID1B*, *CDK6* and *FLT3*/*PAN3* in SEM and OCI-AML3 cells after 24 h MENi (250 nM). The capture viewpoint is shown as a circle at the gene promoters. Menin-bound enhancers in SEM cells are highlighted in yellow.

**Supplementary Figure 4.:**
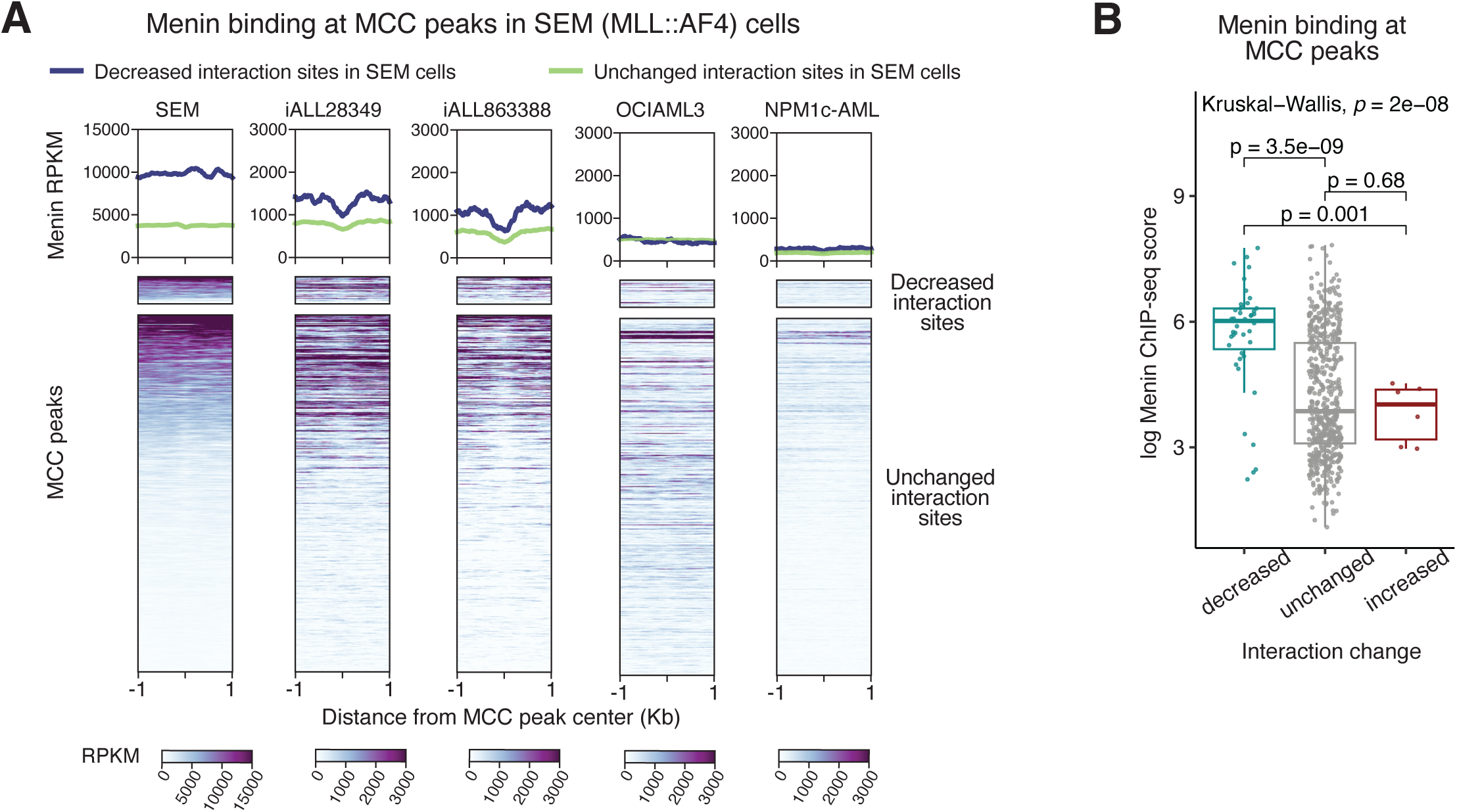
Menin-sensitive enhancers in SEM cells have high levels of Menin binding in SEM and patient samples. (A) Menin binding in cell lines (ChIP-seq in SEM and OCI-AML3 cells) and in primary patient cells (ChIPmentation in iALL28349, iALL863388, and NPM1c-AML) at MCC peaks in SEM cells. Menin signal at MCC peaks with a statistically significant (Mann Whitney U FDR q-value < 0.05, log2 fold change < 0) decrease in unique ligation junctions in SEM cells following MENi (250 nM, 24 h) is shown in dark blue on the metaplots and in the top cluster on the heatmaps. (B) Quantification of Menin ChIP-seq signal at differential MCC peaks in SEM cells.

**Supplementary Figure 5.:**
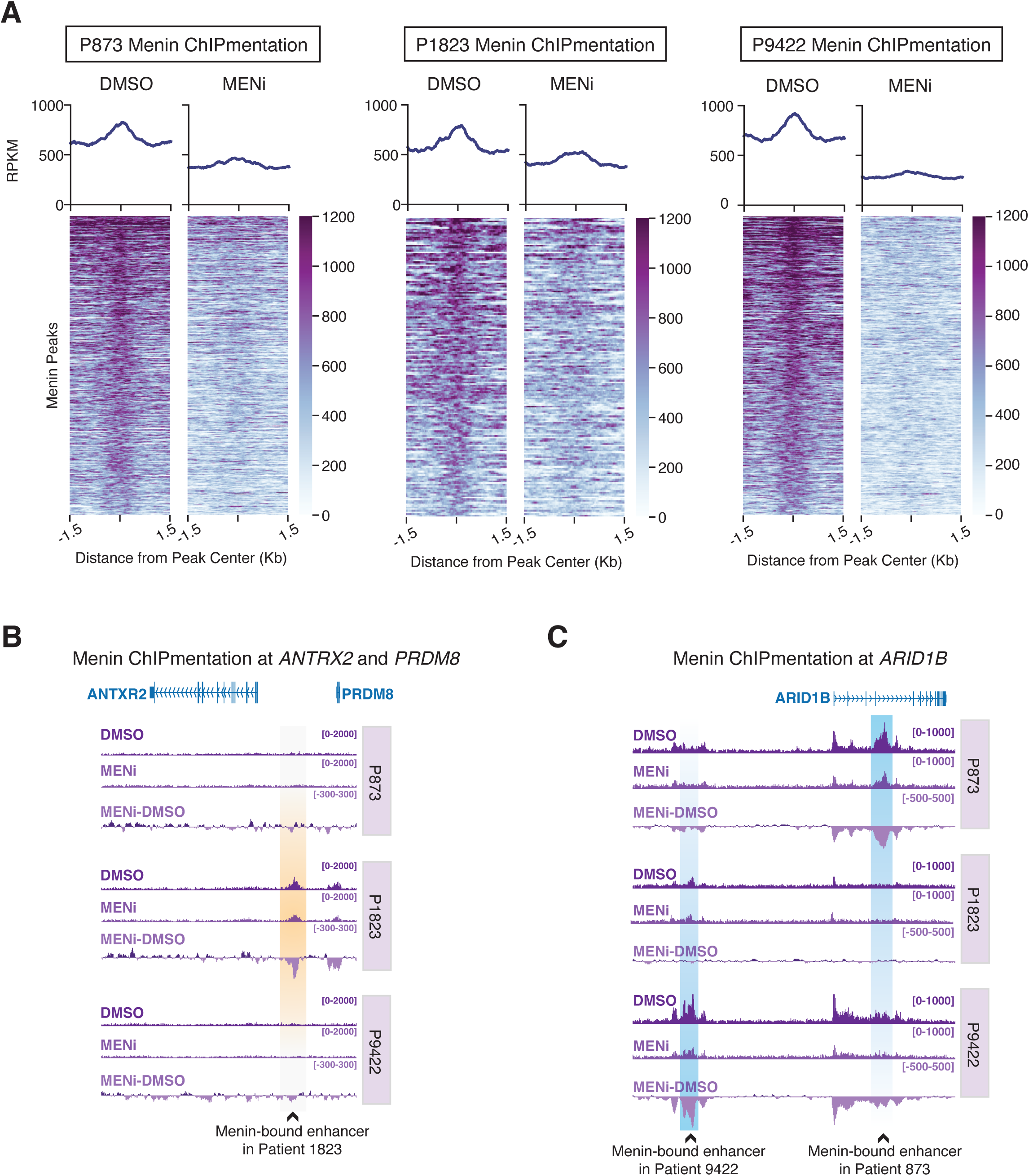
Menin is enriched at enhancers in MLL::AF4 primary patient samples. (A) Menin ChIPmentation at Menin peaks in primary patient cells treated with DMSO or 250 nM MENi for 24 h. (B) Menin ChIPmentation at *ANTRX2* and *PRDM8* in patient cells. (C) Menin ChIPmentation at *ARID1B* in patient cells.

**Supplementary Figure 6:.**
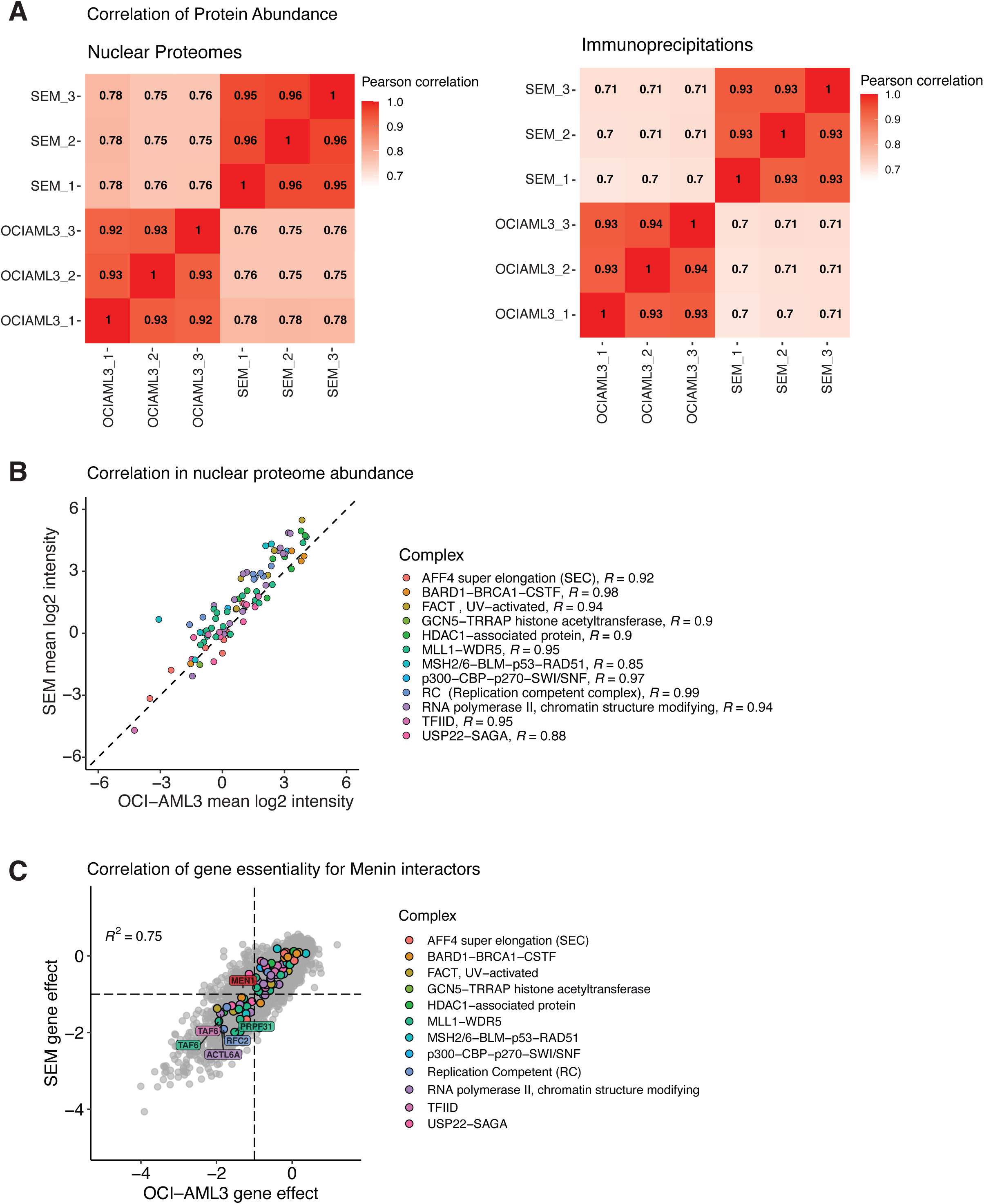
Menin co-IPs have different enrichments of transcription activators despite highly correlated interactomes and nuclear proteomes between cell lines. (A) Pearson correlations between nuclear proteome samples computed on log2-transofrmed and imputed intensities. (B) Correlation in the abundance of differentially enriched complex components in the nuclear proteomes of SEM and OCI-AML3 cells. The grey dashed line represents a slope of 1. Pearson *R* values are shown for each complex. (C) Correlation between the gene effect scores reported in DepMap for SEM and OCI-AML3 CRISPR screens. Proteins within complexes specifically enriched in SEM cells are colored. Genes with an effect score of < -1 in both cell lines are labelled.

## Supplementary Tables

**Supplementary Table 1:** Patient sample characteristics.

| Sample name | % Blasts | Sex | Immuno-phenotype | MLL status | Karyotype | Additional mutations | Protocol |
| --- | --- | --- | --- | --- | --- | --- | --- |
| 9422 | 96 | M | pro-B | t(4;11) |  | NRAS | Interfant-06 |
| 873 | 98 | F | pre-B | t(4;11) | 46,XX,t(2;11;17;4)(q3?1;q23;q25;q21)[15] |  | Interfant-99 |
| 1823 | >90 | F | pro-B | t(1;11) | 46,XX,t(1;11)(p32;q23)[21]/46,XX,[1] |  | Interfant-99 |

**Supplementary Table 2:** Antibodies for chromatin immunoprecipitation.

| Antibody | Reference | Application (per $1 \times 10^7$ cells) |
| --- | --- | --- |
| Menin | Bethyl A300-105A | 2 μg for ChIP, 1 μg for IP |
| MLL N-terminus | Bethyl A300-086A | 2 μg |
| H3K27ac | Diagenode C15410196 | 1.25 μg |
| H3K4me1 | Diagenode C15410194 | 3 μg |
| H3K4me3 | Diagenode C15410003 | 2.6 μg |
| NPM1c | ThermoFisher PA1-46356 | 2 μg |
| Rabbit IgG | Cell Signalling DA1E | 1 ug |

**Supplementary Table 3:** Sources of publicly available sequencing data.

| Dataset | Reference |
| --- | --- |
| OCI-AML3 ATAC-seq | GSM5114291 |
| SEM Menin AutoCUTnTag | GSM5139719 |
| KOPN8 Menin AutoCUTnTag | GSM5139645 |
| RS4;11 Menin AutoCUTnTag | GSM5139689 |
| iALL_28349 Menin ChIPmentation | GSM8657204 |
| iALL_28349 H3K27ac ChIPmentation | GSM8657208 |
| iALL_863388 Menin ChIPmentation | GSM8657184 |
| iALL_863388 H3K27ac ChIPmentation | GSM8657191 |
| MV4;11 Menin ChIP-seq | GSM5858383 |
| MOLM13 Menin ChIP-seq | GSM3635683 |

